# Shifting from correlations to dynamics in the study of human prehistorical resilience to extreme climatic shifts

**DOI:** 10.1101/2023.09.19.558513

**Authors:** Yotam Ben-Oren, Yitzchak Jaffe, Oren Kolodny

## Abstract

In recent years there has been a growing body of research on human resilience to extreme climatic shifts in the past. Most studies focus on comparing archaeological records prior to a perceived climatic shift with those after it, to investigate a causal relationship between the two. Although these comparisons are important, they are limited in their potential to facilitate causal understanding of the factors that determined the human response to climate change. We assert that for such understanding, it is necessary to explicitly consider prior processes that could have made certain populations more resilient to the extreme climatic shift. This assertion calls for a new focus on the cultural and demographic *dynamics* in prehistorical populations, over the generations that *preceded* the climatic shift. In this article, we lay out several mechanisms of cultural evolution that – together with the experienced climatic dynamics prior to extreme climatic shifts – may have determined populations’ abilities to cope with them. This endeavor allows us to outline alternative hypotheses regarding what determined the fate of different human groups. These, in turn, may help direct the collection and analysis of archaeological data and to highlight modalities within it that may be helpful for inference of the mechanisms that determined populations’ resilience to climatic shifts.

## Introduction

The question of how environment and climate affected social development in the past has returned to the forefront of scientific research (Contreras 2016; d’Alpoim Guedes et al. 2016; Degroot et al. 2021; Petraglia et al. 2020; Rick and Sandweiss 2020). The driving force behind this reemergence is the recent availability of new paleoclimate records from across the globe that question old paradigms, coupled with the growing need to understand how societies of different complexities might respond to a changing planet (Bennett et al. 2016; Boivin and

Crowther 2021; Guttmann-Bond 2010; Schiffer et al. 2012). Research has focused on identifying spikes or peaks in climatic and archaeological records to suggest causal links between environmental factors (such as levels of precipitation) and social outcomes (e.g., the collapse of social structures and cultural institutions or a shift in subsistence patterns; see overview in Carelton and Collard 2019). However, these studies usually consider periods between extreme climatic events as though they were largely static phases, and thus rarely go beyond the identification of tipping points in human systems to explain how societies interact with their environments, focusing instead on presumed correlations and seldom considering dynamic processes explicitly (Castree et al. 2014; Holmgren et al. 2016; Kintigh and Ingram 2018).

Prior to exploration of the cultural and demographic processes that determine populations’ resilience to climatic shifts, it is necessary to consider what is meant by population resilience and related terms such as sustainability and robustness, and how different events or dynamics map onto these terms. This is necessary not only for clarity of discussion, but also because this discussion raises some fundamental questions. We will not attempt to provide answers to these questions (nor are we the first to raise them; see overviews in Jaffe et al. 2021; Løvschal 2022; McAnany & Yoffee 2009; Middleton 2017; Redman 2005), but rather to highlight a few of them as topics that must be made clear in discussions of the matter.

The main issue lies with the basic assumptions about resilience, namely that it is the ability of a system to cope with adversity, either by simply weathering/surviving it (aka ‘bouncing back’) or by adapting and changing to accommodate new realities. While the first is often equated with robustness, the latter has been gaining increased traction in many academic circles; though just how much change a system can undergo before it loses its original coherence (and thus can even be seen to have collapsed) is an open matter (Walker 2020). Parsing out what this actually means when dealing with human-environment relationships in the past helps stress this point:

1. Should a population that survives a change of climate be considered resilient if survival was achieved by major cultural changes such as replacement of the social structure or institutions? Alternatively, should a population be viewed as resilient if its culture remained unchanged even if only a small fraction of the population survived? Weiberg (2017), for example, offers a valuable case study, comparing the northeastern Peloponnese and Central Crete. Around 2200 BC, they diverged in their resilience to stress. Weiberg examines adaptive cycles within each region, using millennia of archaeological data to understand population shifts and socio-political changes. While she doesn’t equate population persistence with resilience, her approach considers human continuity, raising questions about how far this concept applies. Is the modern Greek state a current example of resilience spanning millennia since the Neolithic?
2. Is migration a form of population resilience, or population accommodation, of an environmental shift? Or should it be considered a societal collapse (see for example Flohr et al. 2016, contra Weninger et al. 2006)? If the population scattered and each individual or subgroup spread to a different location, or joined other populations, should it be considered a form of resilience, or a failure to accommodate system shocks (van Dommelen 2014; Redman 2005)? How would these be reflected in the archaeological record? For example, one might reasonably assert that successful migration and/or continuation of its material culture should be considered a form of population resilience, however decreased quantities of certain material remains may not readily reflect an opposite scenario, but instead a shift in consumption preferences and social identity negotiations in new homelands (Frachetti 2011; Trabert 2020). Further, in cases in which a population migrated, the archaeological record would usually only show the disappearance or the appearance of a culture, and not both, often leading to a wrong interpretation: a population that successfully migrated to accommodate an environmental change might be viewed as one which had collapsed and disappeared.
3. How should we categorize/perceive cases in which some aspects of society were retained after a climatic shift and others were not? Are some aspects of culture or demography more central for these distinctions? For example, how do we perceive cases in which the subsistence pattern of a society changed significantly, e.g., from farming to herding, while social institutions such as religion or government remained relatively unchanged — an often cited outcome of the so-called ∼4k/4.2kcalbp climate anomaly in places like northern China where populations are believed to have adapted by shifting from sedentary agriculture to increased dependence on more nomadic forms of pastoralism for economic survival and with it changing their sociopolitical makeup (e.g. An et al 2005; Liu and Fang 2012 and see Jaffe and Hein 2020).
4. Is occupational continuity of a site by the same population evidence for resilience? Should it matter if the identity of the human population changed, and if so – what kind of change “matters”? Consider different scenarios as examples: (A) The same genetic lineages persist in the same locality, but take on different material or ideological characteristics; (A1) borrowed/incorporated from elsewhere, or (A2) developed locally. (B) Many social aspects – subsistence pattern, religion, and/or political structure – persist, but the people change; the population, gradually or suddenly, is replaced by different genetic lineages / a different ethnic group (e.g., Posth et al. 2018). (C) Both culture and ethnicity/genetics change, but the site is continuously occupied. (D) A certain part of society persists, e.g., only the political/economic elite or only the agrarian workers, or only the urban dwellers, and others perish or migrate. The current surge in aDNA studies serves as a prime illustration of these inquiries, with scholars engaging in discussions about the true implications of genetic marker variations throughout history and geography; Further complicating matters is the perspective of those interpreting the data and their reliance on specific theoretical frameworks, which shape how material remains are perceived to reflect or link with the above-mentioned social elements (Booth et al. 2019; Bruck 2021; Eisnmann et al 2018; Reich 2018)
5. Is retention of population size an important factor for considering whether a population was resilient to a climatic shift? How much of a change, namely demographic decline, constitutes a significant change? Discussions on the influence of environmental factors on Maya societies and political structures have squarely addressed this dilemma: While the upper strata of society, along with their associated political institutions, experienced a collapse, the population centers endured. The critical question remains: To what extent can we genuinely attribute this event to the impacts of climate change (Hogarth et al. 2017; McAnany & Yoffee 2009; Turner and Sablof 2012)?
6. What is considered a climatic shift, enough to impact human populations, is itself a much-debated question (Degroot et al. 2021; Jaffe et al. 2020; McPhillips et al. 2018; Stoot et al. 2016): What is climate change and what is climate variation? Is there a change “for good” (e.g., more rain) or “for bad” (e.g., drought), or are all climatic shifts equal, and what matters is the absolute difference in values of defined variables of interest? Some variables that come to mind may be temperature and rainfall, but what measure of these, and on what time scales? Daily/monthly/yearly mean? Most extreme values? Frequency of extreme values? Frequency of crossing certain thresholds? Arbitrary thresholds, or ones that need to be linked to practices, e.g., the mean annual amount of 200mm rain considered necessary for wheat farming (see in Rosen and Weiner 1994).
7. What is the essence and what is the scale of environmental change that we should study? Some possible examples: changes in ecological carrying capacity (bioproductivity); ecosystem changes such as the ratio of wooded space to savannah; change in the environment that alters the appropriateness of cultural practices to the ecosystem dynamics (e.g. change in frequency of flash floods, mudslides, or wildfires, and the suitability of housing style to accommodate these).

How is cultural or demographic change defined? Must it be sudden, and clearly co-localized in time and space with the climatic shift, or should gradual change, following the shift or cooccurring with it, be considered a form of accommodation/persistence (Diamond 2005; Faulseit, 2016; Haldon et al. 2016; Izdebskiet al. 2016)? What are the variables/thresholds that need be considered? If variance in any of these was very high prior to the climatic shift, and the variable values are within this variation change, should it be viewed as a form of accommodation, or as persistence without cultural change? This may lead to a situation in which by one (technical) definition of resilience, cultures that have more variance in the variables of interest would be considered more resilient (Middelton 2017). Finally, can collapse be intentional? Hopi traditions tell of cases where early communities, which were thriving economically, abandoned their settlements and existing social constellations upon realizing that they had fallen into a state of ‘corrupted life’. In their perception, in such cases the community must move on, destroy the existing group in order to find itself anew – the only way to survive is to intentionally commit social collapse (Fowles 2016).

We leave these questions open: we do not think any of them have a correct or incorrect answer. Rather, these are questions that need to be considered and kept in mind when discussing human responses to climatic shifts and their exploration. This is essential both for facilitating effective communication, and for informed exploration of the aspects of data or dynamics that might best inform researchers regarding what constitutes pertinent questions or variables. Crucial for our discussion here is that not only has the vagueness and open-ended use of terms such as “resilience” hampered in-depth understanding of past human-environment dynamics, but when this issue is highlighted (and numerous papers have done precisely that), they have rarely, if ever, addressed the issue of how repeated longer-term dynamics influence and shape human reactions to climate change and their various outcomes.

In what follows, we lay out several alternative dynamics of cultural evolution that precede a major climatic shift, and we consider two climate-related scenarios: one in which climate is stable prior to the major shift, and a second in which the climate goes through repeated climatic shifts of an intermediate amplitude prior to the major shift. For simplicity, these shifts are short, with a duration on the order of a single time unit, and are thus also referred to as “climatic events”. Here, for simplicity, these events are always considered negative in their direct effect on the population, reflected in a decrease in population size.

### The model and the explored scenarios

We propose a dynamic, spatially explicit model to simulate how experiencing low-magnitude environmental events affects meta-populations’ resilience to extreme climatic disasters, exploring a range of mechanisms of cultural evolution. These mechanisms may determine a population’s response to environmental change. The model consists of a grid of 10X10 populations. Metapopulation dynamics were simulated in two phases. In the first phase, lasting 500 time-units, the population experienced environmental events at different probabilities (0 to 0.5 per time unit, in different simulations). Next, the metapopulation experienced an extreme climatic disaster, similar in nature to the previous events, but greater in magnitude, after which the metapopulation dynamics were simulated for a further 100 time units without environmental events, to examine recovery. In the presented results, all populations start from the same initial population size, *N*=1000 individuals, and are assigned the same carrying capacity, *K*=2000, which limits their growth. In each time unit in which a climatic event does not take place, each population grows following the commonly used discrete logistic growth function (N_t+1_ = N_t_ + N_t_ × r × (K – N_t_); Verhulst 1838, Otto and Day 2007), where *r* signifies the growth rate and was set to *0*.*1*. After calculating the expected population size using the growth function, to add stochasticity, we then draw the actual population size from a normal distribution, where the mean is the expected population size, and the standard deviation is 5% of that size. At the beginning of each simulation, each population is assigned an environmental-event-fitness value, *f*. A population’s fitness value determines its ability to accommodate environmental shifts, i.e. it reflects some population-level know-how that facilitates survival in the face of adverse climatic events. This value can initially be either uniform in all populations across the grid or a randomly drawn number between 0 and 1 for each population, depending on the scenario explored. Every environmental event has a magnitude of *m*, and *f* dictates the fraction of this maximal magnitude that is experienced by the population. Thus, when an environmental event happens, each population’s size is reduced, following this equation:

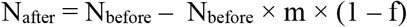

For example, if the population size before the event, *N*_*before*_, is 100, and the magnitude of the event is 1, for *f* = 0, the population size (*N*_*after*_) will go down to 0, for *f* = 0.5, it will reduce to 50, and for *f* = 1, it will not change. Environmental events’ *m* was drawn **per population** from a normal distribution, with a mean of 2 and a standard deviation of 1. Disaster *m* was set to 5. Each case in which a population’s size decreases below *N*=10 is considered a population collapse, and the population is deleted from the metapopulation grid, i.e. for the sake of simplicity in the current discussion, we use the term “collapse” to describe a case in which the population ceases to exist. Because different populations have different *f* values and different census sizes, and because the exact magnitude of each event is drawn per population, different populations may have different fates following an environmental event – some may collapse, while others may go virtually unharmed.

Whenever a population goes extinct, its location in the grid can be inherited by individuals from a surviving neighboring population. If available, a neighboring population at carrying capacity is randomly chosen to send a reproduction surplus of 10 individuals to establish a new population in the vacant location, inheriting its parent population’s fitness.

### Explored Mechanisms

In order to elucidate potential mechanisms and delve into their implications for influencing a population’s response to a profound climate shift, we conducted individual examinations of the following hypothesized mechanisms. It is important to note that these mechanisms are not strictly isolated from one another; they may operate concurrently. Investigating their combined impact and the interplay between them represents a valuable pursuit, extending beyond the scope of our current study.

#### 1) Necessity is the mother of invention

In this example, we explore a scenario in which environmental events lead to cultural change, resulting in a change in *f*. There can be many reasons for such a trend. Some scholars have suggested, for example, that cultural innovation could be a product of necessity (e.g. Boserup, 1975). Moreover, studies in behavioral ecology suggest that behavioral conservatism would be generally preferred evolutionarily, with one of the rare exceptions being desperate situations where conservative behavior cannot guarantee survival (Kacelnik and El Mouden 2013; Greenbaum et al. 2019). Similarly, there is genetic evidence that similarly shows an increased mutational rate under stress, which could be also a case of adaptive “exploration” (e.g., see Wei et al. 2022.) Alternatively, increased mutation rate/cultural change could be a result of difficulty in conserving the existing genetic/cultural composition under stress. Either way, these examples suggest that populations may be less likely to stick to their known cultural practices under stress. Depending on the specific mechanism explored, cultural change following environmental events can be either directional (=improvement in facing environmental challenges) or random (as in genetic mutations, which can be positive or negative). Here, we chose to explore the more conservative option, i.e., that the directionality of change in *f* is random, in order to demonstrate that even when cultural change is not necessarily positive, environmental events are expected to increase the metapopulation fitness.

In this scenario, all populations are assigned the same *f*, 0.7, at the beginning of the simulation. Here, we assume that *f* is, in itself, a product of environmental change. When an environmental event occurs, *f* “mutates”, drawing from a normal distribution where the mean is the previous *f* and the standard deviation is 0.05.

#### 2) Selection on existing variance

Unlike in the previous scenario, here we explore the possibility – perhaps the more parsimonious option – that any change in the metapopulation’s fitness is not a result of cultural innovation or exploration but rather of selection on preexisting cultural practices. In other words, here we assume that when the metapopulation faces environmental events, populations with lower *f* are removed and replaced by more successful populations, a process which ultimately increases the average fitness in the metapopulation.

In this example, at the beginning of the simulation, each population is assigned a random *f* between 0 and 1, which does not change as long as the population survives. Any change in the metapopulation’s average fitness is therefore a result of selection among populations and repeated replacement of populations that went extinct; these, on average, tend to be those with lower *f* values.

#### 3) Experience-based cultural transmission

In this version of the model, we assume cultural knowledge relevant to coping with environmental events is transmitted by experience and is likely to be lost in the absence of environmental events. One can imagine that the knowledge relevant to surviving droughts can be lost if there is no need for it for several generations; or that a costly fortification system, built to protect a village from floods, will not be maintained over time unless it is used occasionally.

In this scenario, all populations begin with the same high *f* value, 0.95, and it can decay over time if the population does not practice dealing with environmental events. The probability of cultural loss, *P*_*loss*_, for which we used the value 10^-4^, is multiplied by the number of time units since the last environmental event, meaning that the longer the knowledge is not used, the more likely some of it will be lost. In each cultural loss event, the population loses 10% of its current *f*.

#### 4) Demography-dependent rate of invention

A common hypothesis in the field of cultural evolution is that cultural innovation is positively correlated with demography (Powell et al. 2009, Kolodny et al. 2015). One reason for this is that if every individual in a population has a given probability of inventing a new cultural trait, the larger the population, the more inventions are expected to occur. Alternatively, some suggest that larger populations are able to sustain more specialized guilds/institutions, which in turn can increase their overall cultural repertoire (Ben-Oren et al. 2023). Populations that experience frequent environmental events where their size is reduced are, on average, smaller than populations that experience environmental stability and may also be potentially less successful in maintaining many cultural specializations (Ben-Oren et al. 2023). Therefore, frequent environmental events may decrease the accumulation of cultural adaptations.

In this scenario, population fitness is accumulated via cultural inventions. All populations begin with the same value of *f*, 0.7, and inventions occur with a probability *P*_*inv*_, set by default to 0.001, which is multiplied by the population’s size (assuming each individual has the same probability of invention, similar to Kolodny et al. 2015).

#### 5) Demography-dependent rate of cultural loss

The ability to maintain the cultural repertoire, which in our case translates to the maintenance of fitness to cope with environmental events, may also be positively associated with population size. For example, the more individuals know a certain cultural trait, the less likely it could be for it to be lost by chance (Kolodny et al. 2015). Similarly, the more individuals who know a certain trait, the more likely a good demonstrator will be found to transmit the cultural knowledge (Henrich 2004). Thus, the probability of cultural loss may be greater when population size is smaller.

In this scenario, all populations begin with the same high fitness, 0.95, and loss events occur in a probability *P*_*loss*_, set by default to 0.05, which is multiplied by the fraction of individuals “missing” between the census size and the carrying capacity (*P*_*loss*_ × *(K – N) / K*). The idea behind this is that the higher the proportion of empty positions in a population, the higher the probability for cultural loss. In each cultural loss event, the population loses 10% of its current *f*.

## Results

For each scenario, the overall number of individuals in the metapopulation was monitored throughout the simulation (i.e., before and after the disaster) for different probabilities of environmental events per time unit, ranging between 0 and 0.5. Here, we define metapopulation resilience as maintenance of a high number of individuals in the metapopulation following an environmental event (while other possible definitions are discussed in the Introduction and Discussion). In three of the explored scenarios (“Necessity is the mother of invention”, “Selection on existing variance”, and “Experience-based cultural transmission”), intermediate frequency of environmental events seems to increase metapopulation resilience, while in the remaining two (“Demography-dependent rate of innovation” and “Demography-dependent rate of cultural loss”) they decrease resilience.

1. Necessity is the mother of invention
2. Selection on existing variance
3. Experience-based cultural transmission
4. Demography-dependent rate of invention
5. Demography-dependent rate of cultural loss

## Discussion

Our explorations show that very different processes – modelled within the same framework to facilitate comparison and explicit layout of underlying assumptions – may lead to similar outcomes in terms of demography. For example, a population’s persistence in the face of a major environmental crisis might be due to increased cultural innovation when climate change begins to occur and conditions become unstable, or it might be the result of recurring smallscale climatic events that induced natural selection among cultural variants that existed in the metapopulation, favoring variants that provided resilience to severe climatic events early on. Presenting diverse scenarios within a unified framework that can either bolster or impede resilience in the face of climate-related challenges is essential for formulating hypotheses regarding the factors that may have influenced the development of past populations.

This approach may also enable us to emphasize overlooked aspects of empirical data that can differentiate between various scenarios, even when their anticipated demographic outcomes— often the primary focus of population resilience studies in archaeology—appear similar. For example, we show in scenarios 1 and 2 (“necessity is the mother of invention” and “selection on existing variance”) that the expected demographic outcomes are similar, but the expected trend in cultural variation is different (see Figures 1 and 2). While in scenario 2, variance is only reduced following environmental events, in scenario 1, we see generation of new variance following every event. Likewise, in both scenarios 4 and 5 (“demography-dependent rate of invention” and “demography-dependent rate of cultural loss”) environmental stability seems to promote resilience, but in scenario 4 the overall trend is an increase in resilience, while in scenario 5 the trend is opposite. In addition, in scenario 4 even populations which experience frequent environmental events accumulate inventions gradually (albeit at a slower pace), which means that if they manage to survive long enough, they may eventually become resilient to environmental stress. Similarly, in both scenarios 3 and 5 (“Experience-based cultural transmission” and “demography-dependent rate of cultural loss”), there is an increase in *f* immediately after environmental events, as a result of selection against populations with low fitness, followed by a reduction in *f* until the next event. However, in scenario 3 the reduction is a product of limited experience in dealing with environmental events, and it thus accelerates as more time passes since the last event, while in scenario 5, the reduction results from reduced population numbers, which makes it more likely that decline in *f* would occur right after environmental events and before the population has recovered. In other words, while in scenario 3 the reduction in *f* accelerates as time passes since the last event, in scenario 5 it slows down. Such realizations may direct future archaeological efforts to better interpret past population dynamics.

**Figure 1.**
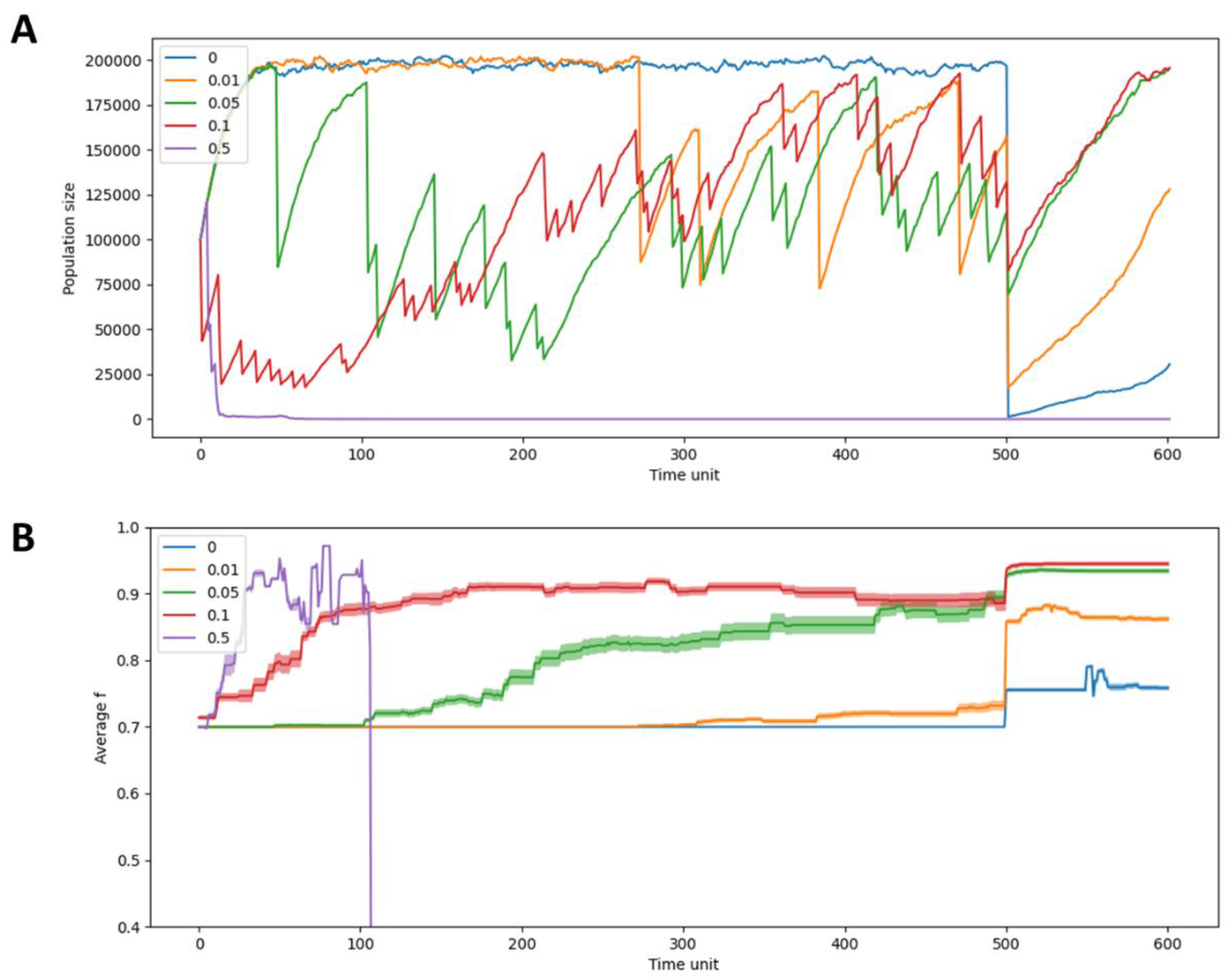
Cultural change following environmental events increases resilience in populations experiencing intermediate rates of events. Panel (A) shows the total number of individuals in metapopulations experiencing different frequencies of environmental events throughout the simulation. Panel (B) shows the average *f* values for the same metapopulations, with variance shown by the shaded areas. In this example, environmental events lead to the generation of new variance in *f*, as well as an increase in the average *f*. We find that in this scenario, experiencing extremely high frequencies of environmental events (0.5 per time unit) led to the collapse of the metapopulations even prior to the environmental disaster. However, intermediate frequencies of environmental events led to increased demographic resilience of the metapopulations, both in relative and absolute terms (with the metapopulation experiencing environmental events with a probability of 0.1 showing the highest resilience). Notably, a gradual adaptation of metapopulations experiencing environmental events is evident before the environmental disaster – both in the increase in *f*, and in the gradual population growth throughout the phase preceding the disaster. In this example, intermediate frequency of environmental events is also associated with the highest variance in *f* values, as environmental events both increase variance by introducing cultural change and reduce it by selecting against populations with low *f* values.

**Figure 2.**
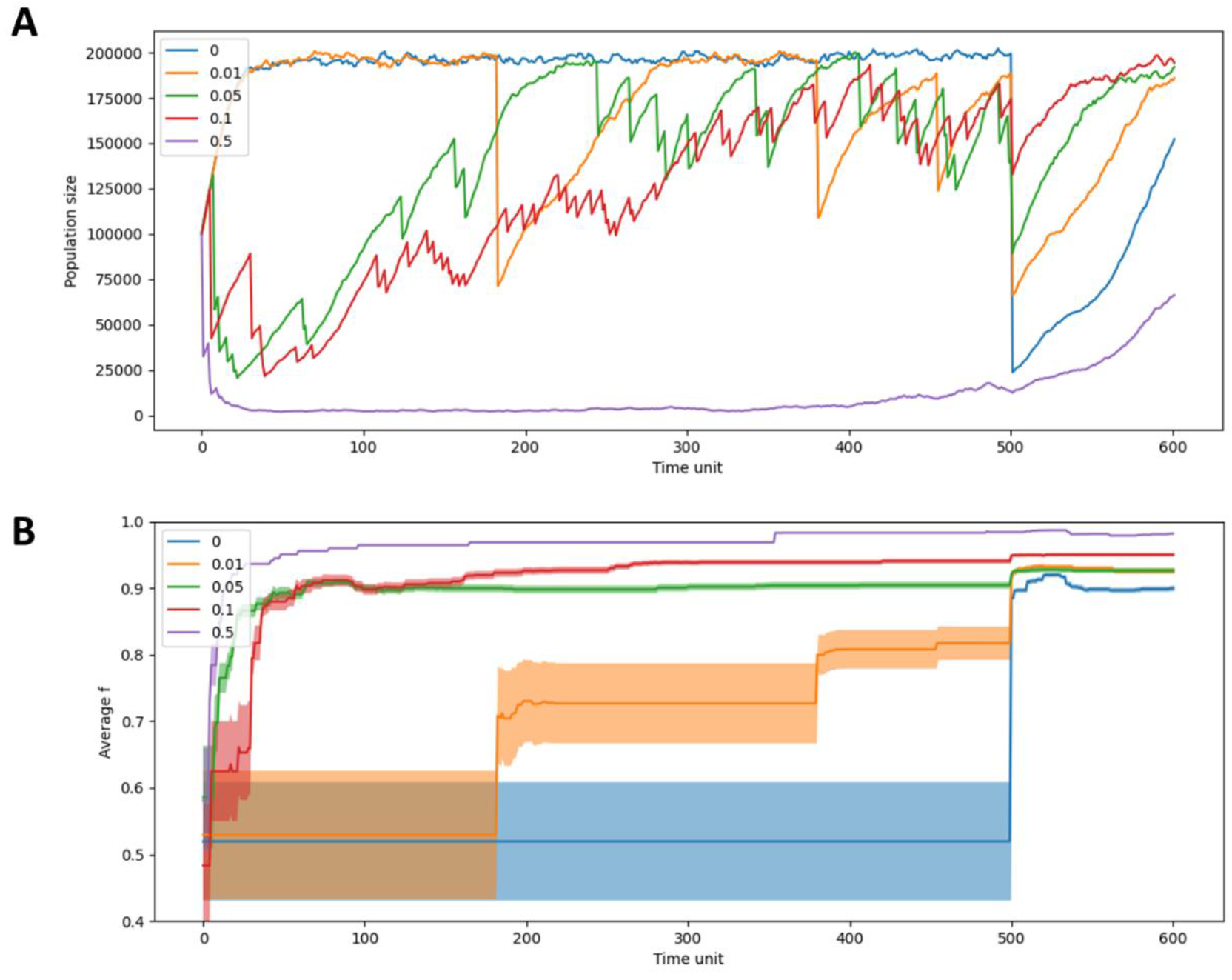
Selection against populations with low *f* increases resilience in populations experiencing intermediate rates of environmental events. Panel (A) shows the total number of individuals in metapopulations experiencing different frequencies of environmental events throughout the simulation. Panel (B) shows the average *f* values for the same meta population, with variance shown in the shaded areas. In this example, the metapopulation has a high variance in *f* in the beginning of the simulation, which is gradually reduced following environmental events, as populations are removed and replaced. At the same time, each environmental event is followed by an increase in the average *f* (See, for example in the orange graphs). Because we assume populations with high *f* exist from the beginning of the simulation, regardless of the environmental events that follow, in all cases the metapopulation seems to survive the environmental disaster. However, intermediate frequency of environmental events seems to reach the highest population sizes after the environmental disaster (with populations experiencing environmental events in a probability of 0.1 showing the highest resilience). Similar to the previous scenario, here we also see that metapopulations that experience intermediate frequency of environmental events show a gradual increase in resilience (as marked by population number increase, and increase in the average *f*) before the environmental disaster.

**Figure 3A.**
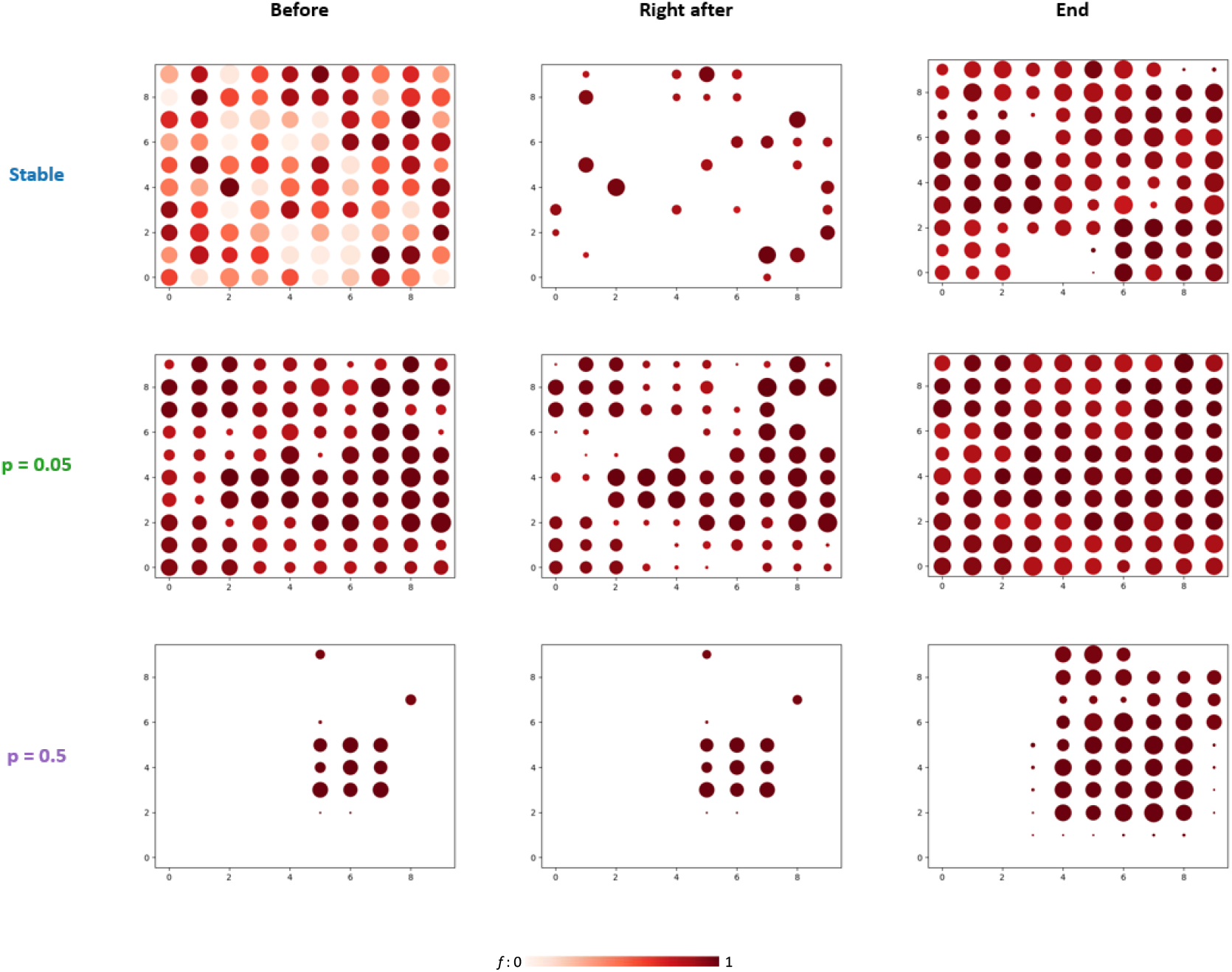
Metapopulation maps before the environmental disaster, immediately after it, and at the end of the simulation for metapopulations experiencing environmental stability (no environmental events), intermediate probability of environmental events. (0.05 per time unit), **and high probability of environmental events** (0.5 per time unit). In Figure 3A, every population is colored according to its *f* value (with higher values represented in darker colors). In Figure 3B, every population is colored according to its lineage – meaning that at the beginning of the simulation, each population is assigned a color, and later, if members of this population establish a new population, it is assigned the same color. Population size is represented by circle size, with the largest circles representing populations at carrying capacity (*N* = 2000). Until the environmental disaster, the highest diversity of lineages and *f* values, as well as the highest metapopulation size, are maintained in the metapopulations experiencing environmental stability. When the environmental disaster occurs, most of the populations collapse since many of them have low *f* values. In populations experiencing extremely high frequency of environmental events, on the other hand, diversity before the environmental disaster is minimized. In terms of lineage, out of 100 lineages at the beginning of the simulation, only 4 survived until the disaster (two of which are represented in very similar colors), all with limited spread. In terms of diversity in *f* values (as can also be observed in Figure 2), only populations with *f* values that are very close to 1 survived. However, when the environmental disaster occurs, it has a minimal effect: all populations survive, and the demographic reduction is minor. In populations experiencing intermediate frequency of environmental events, these are frequent enough to allow for selection in favor of higher *f* values, but also rare enough to allow for demographic recovery and survival of multiple lineages. Thus, while in such metapopulations, the number of individuals might not be as high as in metapopulations experiencing environmental stability, and their average *f* may not be as high as in metapopulations experiencing frequent environmental events, the number of populations that survive the disaster is the highest.

**Figure 3B.**
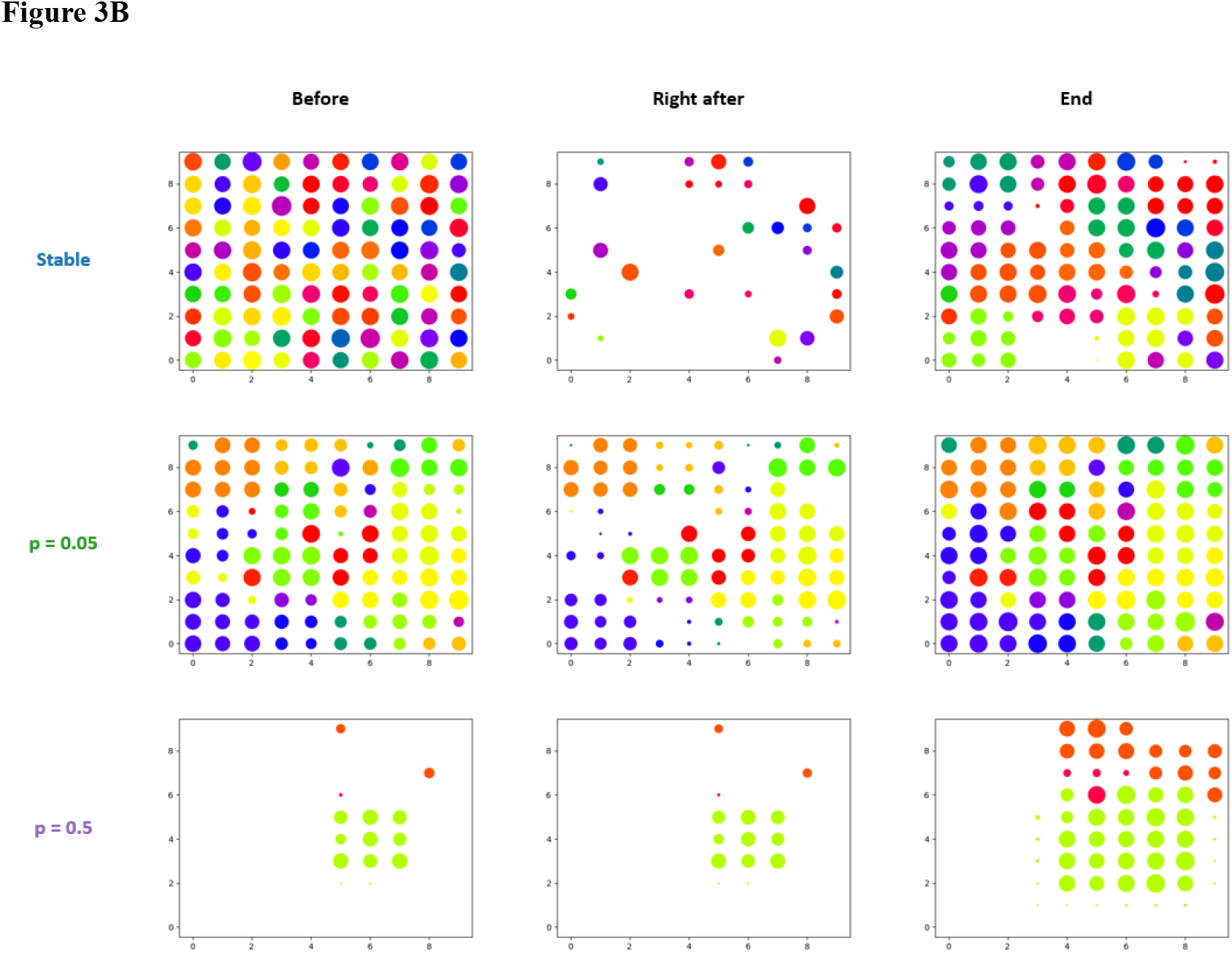

**Figure 4.**
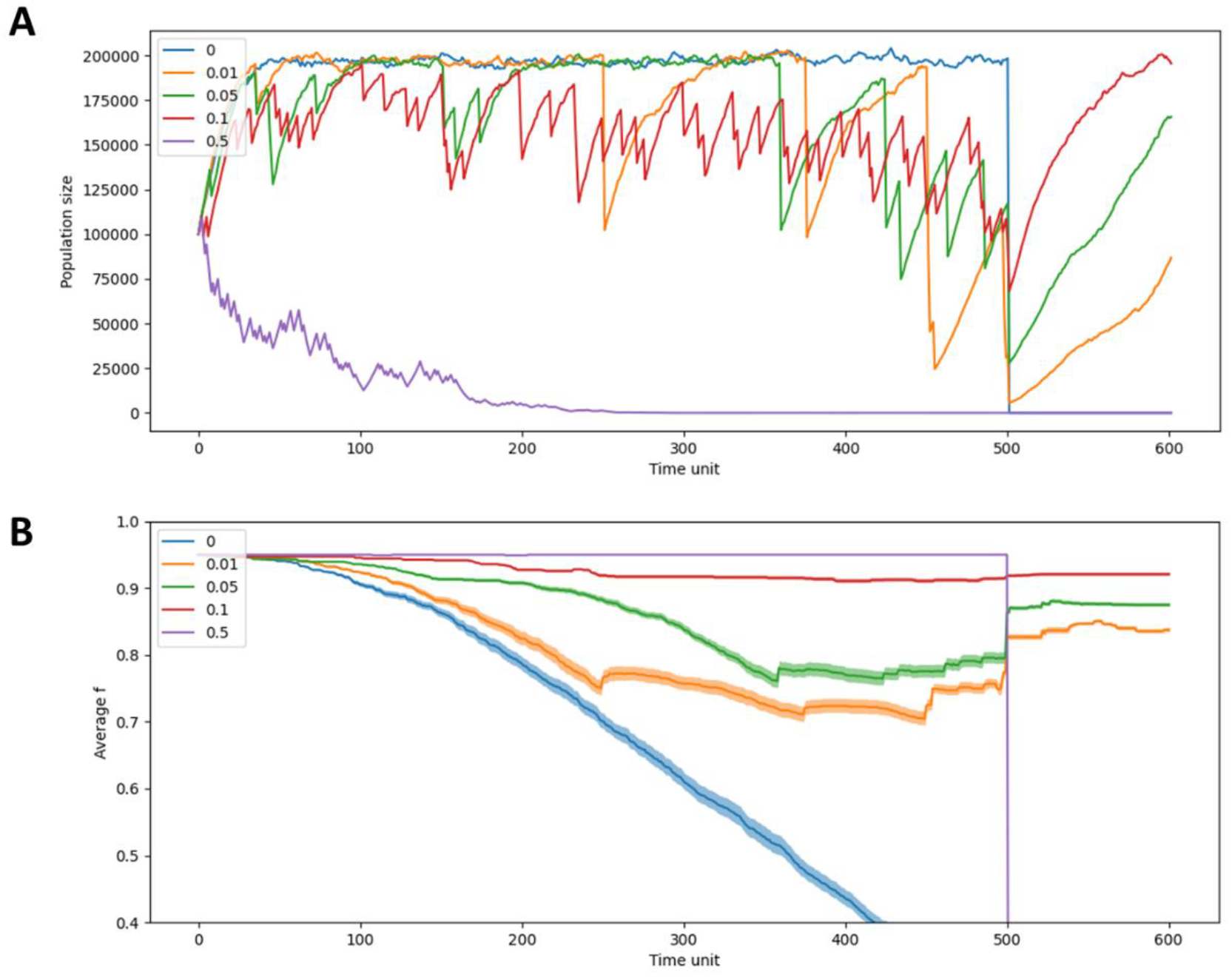
When the cultural knowledge needed to cope with environmental change is maintained through experience, intermediate frequency of environmental events contributes to metapopulation resilience. Panel (A) shows the total number of individuals in metapopulations experiencing different frequencies of environmental events throughout the simulation. Panel (B) shows the average *f* values for the same meta population, with variance shown in the shaded areas. In this scenario, all populations had high *f* values (0.95) at the beginning of the simulations, but these were likely to be reduced in the absence of environmental events. Like the dynamics in the previous scenarios, we find that metapopulations experiencing intermediate frequency of environmental events remain the largest following an environmental disaster (with populations experiencing eventss with a probability of 0.1 showing the highest resilience). However, unlike the previous scenarios, here *f* values gradually decrease, which means the effect of environmental events becomes more severe over time. We find that after environmental evets the average *f* initially increases, as a product of selection, and then starts to decrease. The decrease is initially slow but gradually accelerates, until the next environmental event occurs. That is because the probability of cultural loss increases, the longer the time since the last environmental event.

**Figure 5.**
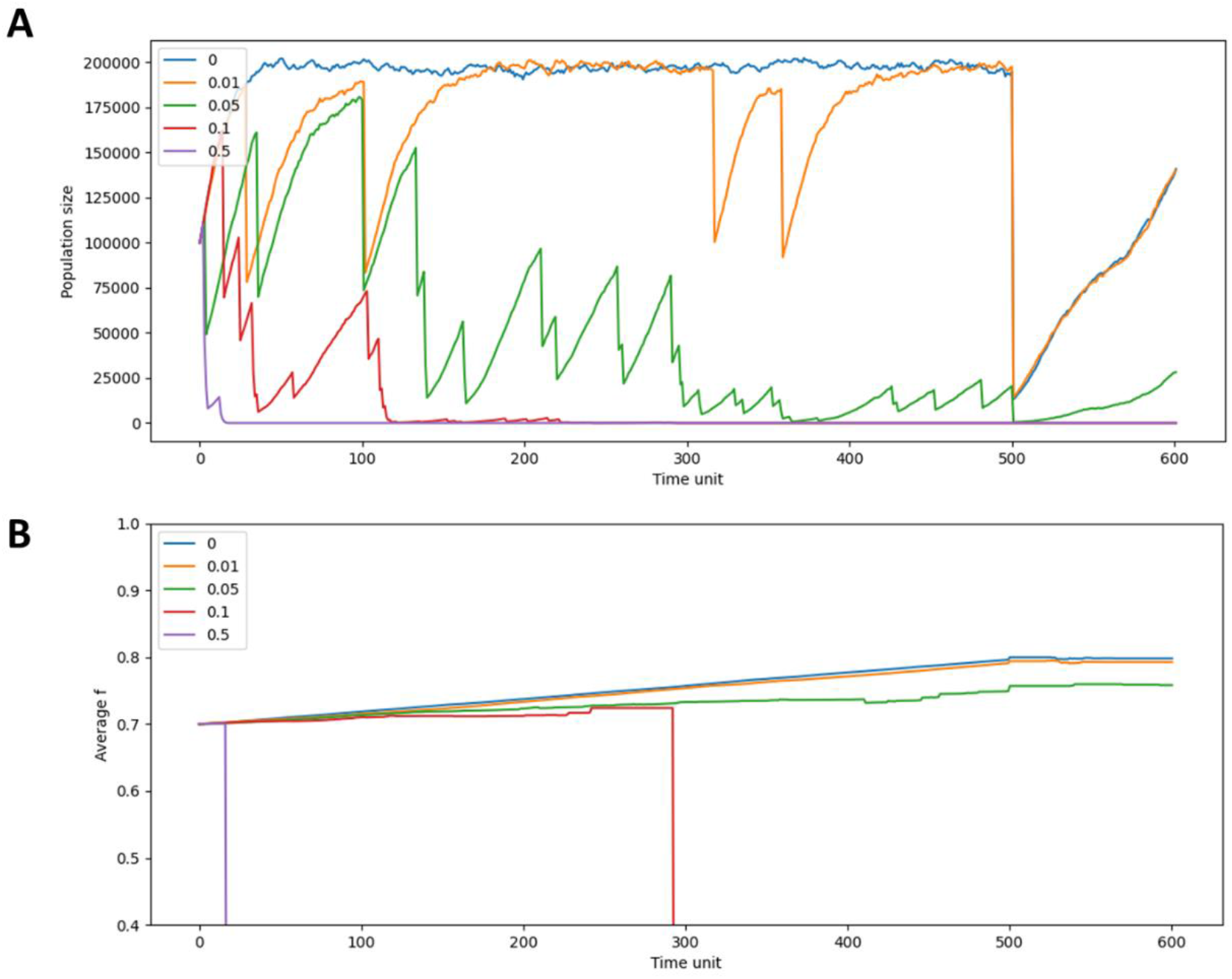
When the rate of accumulation of cultural knowledge is dependent on population size, environmental stability contributes to population resilience to environmental events. Panel (A) shows the total number of individuals in metapopulations experiencing different frequencies of environmental events throughout the simulation. Panel (B) shows the average *f* values for the same meta population, with variance shown in the shaded areas. In this scenario, all populations start with an intermediate *f* value (0.7), which can increase by cultural inventions, appearing in a probability proportional to the population size. We find that under these assumptions, metapopulations that experience frequent environmental events may collapse even before the environmental disaster and that those who do survive until then, are more dramatically affected by it than metapopulations that experience environmental stability. Because populations that experience environmental events are on average smaller, the rate of accumulation of *f* is smaller, which makes them more fragile, when the disaster arrives.

**Figure 6.**
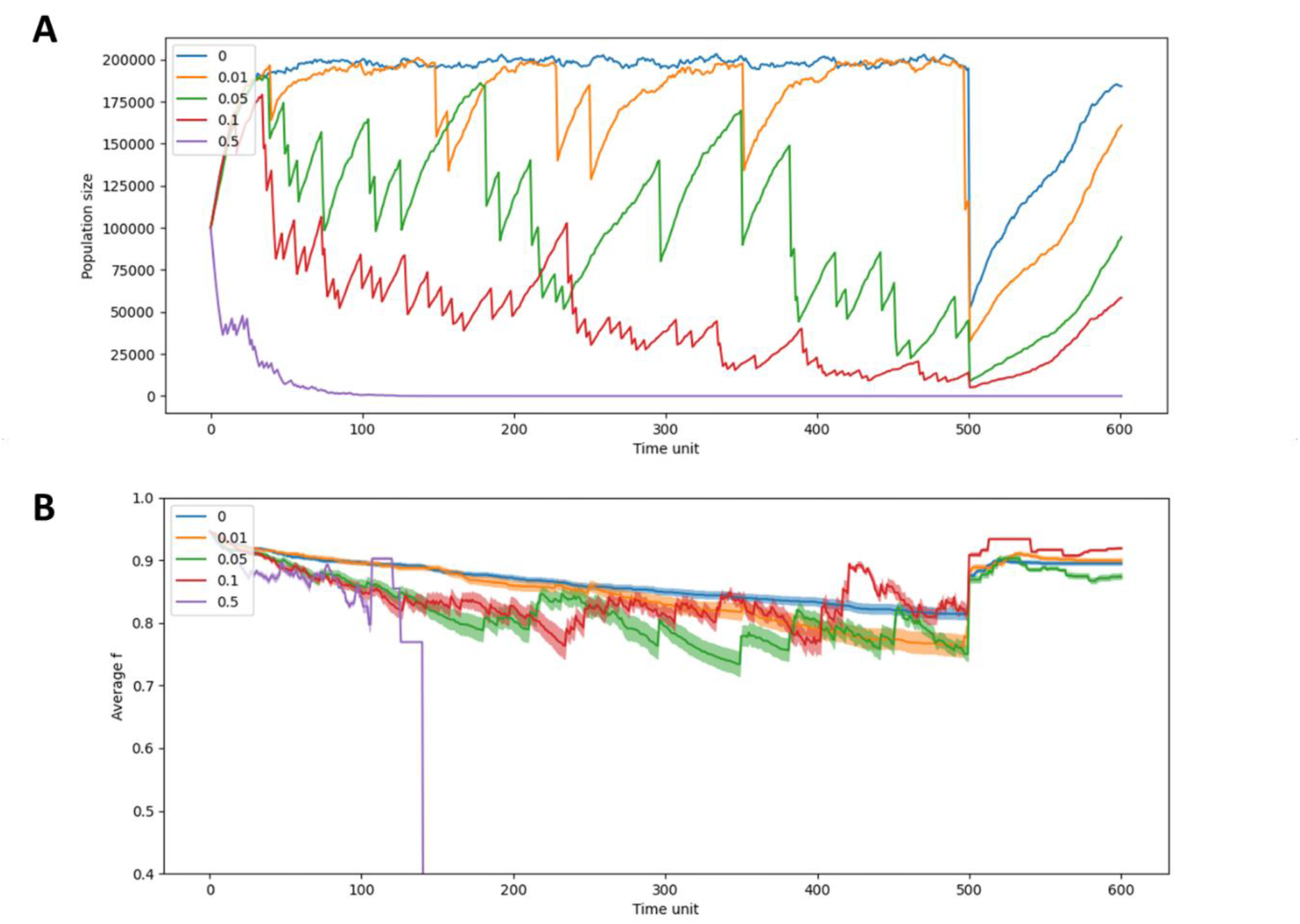
When the probability of loss of cultural knowledge is dependent on population size, environmental stability contributes to population resilience to environmental events. Panel (A) shows the total number of individuals in metapopulations experiencing different frequencies of environmental events throughout the simulation. Panel (B) shows the average *f* values for the same meta population, with variance shown in the shaded areas. In this scenario, all populations are initially assigned a high *f* value (0.95), which may decrease, due to loss of relevant cultural knowledge. We assume, in this case, that smaller populations have a higher probability of cultural loss, and find that under these assumptions, frequent environmental events lead to cultural loss, and thus to reduced resilience to environmental disasters. Since average *f* gradually decreases, here we find that environmental events lead to greater reductions in metapopulation size over time. Also, unlike in the other cultural loss scenario (“Experience-based cultural transmission”), here we see that after environmental events average *f* drops the fastest (as then population sizes are the smallest) and then the pace of reduction in *f* slows down (as populations grow).

Returning to our original set of questions regarding assumptions in the existing literature on human-environment dynamics in the past: our study does not answer these questions; rather, it adds qualitatively different questions to this list, as it seeks to engage with often overlooked aspects of the long-term relationships between humans and their environments (and see Kintigh et al 2019; Haldon et al. 2018; Hussain and Riede 2020). The above models exemplify the intricate relationships between changing environments and social coping mechanisms, highlighting the problem, for simplicity, by treating individual climatic events as inherently catastrophic and equally disastrous for all societies across time and space. However, this exploration shows that although each event has a direct negative effect, recurring events may have in some cases a positive impact on a population’s long-term resilience when faced eventually with an extreme event. This exemplifies that to understand the impact of climate on a society, it is imperative to consider long-term processes: previous environmental dynamics, population sizes, and the society’s capacity to address various climatic anomalies.

The results of this study suggest that populations experiencing at least some rate of environmental crises might experience less shock when large-scale events hit. Notably, throughout all five scenarios, environmental events allowed for selection in favor of populations with high *f*, meaning that regardless of the specific consequences of these events on populations (e.g., if they make cultural loss more likely) they have some positive consequences for the metapopulation’s long-term resilience. The models we tested are only initial steps in addressing these issues. The question of how frequent and intense these events can be (as in what is the ‘sweet spot’ between too much and too little stress) is one which archaeologists, working together with paleoclimatologists, may be able to explore empirically. An additional avenue of exploration is how stochasticity plays into the formulation of environmental fitness (Burke et al. 2021; Kennet and Marwan 2015). Would a system that experiences random small events fare better or worse than one that undergoes more intense or more frequent events, but in a more predictable manner (e.g. seasonal flooding, monsoon rains, dry seasons etc.)? Similarly, can stochasticity and lack of predictability in intensity and timing turn out to be beneficial for those coping with climate change?

Settlement persistence has become an emerging trend in archaeological literature focused on understand the contributing factors to longevity in territorial occupation (Craford et al. 2023; Feinman and Neitzel 2023; Orton et al. 2020; Smith et al. 2021), with some finding urban systems better positioned to tackle changing climates given their institutional reserves and increased social complexity (see in Lawrnace et al. 2021). We highlight an additional contributing element to population survival, focusing on the long-term interplay between recurring environmental shifts and population size. Archaeologists and historians can address environmental volatility to assess how aspects of urban resilience, contra smaller villages for example, may lead to continued persistence over time.

The scenarios we explored in our research provide examples of different categories of effects that environmental events may have on population resilience. There is, however, ample opportunity for further exploration. For example, while, for simplicity, we did not account for the effects of inter-population contact on resilience, we appreciate this is also likely to affect their cultural repertoire (Ben-Oren et al. 2023) and their demographics. Similarly, we assumed that populations’ *f* affects them only during environmental events while it may also affect them in periods of environmental stability (e.g., high *f* could also allow for faster recovery from environmental events or, contrarily, there could be a trade-off between high *f* and growth rate under stability). Additionally, besides the five scenarios studied here, many more can be explored. For example, environmental events may have a range of effects on populations’ social structure, which could, in turn, affect their resilience to disasters. Alternatively, populations may have material reserves that could interact in different ways with population size, resilience, and cultural accumulation. Furthermore, in our models, we explored general scenarios, which can result from different specific mechanisms. For example, a positive association between the number of individuals in a population and the rate of cultural innovations can emerge because all individuals have a similar probability of coming up with inventions, and thus, there is a linear relationship between population size and the rate of inventions in the population. Conversely, this can also be the result of larger populations allowing for more cultural specializations/institutions, which may create a non-linear relationship in the same direction. While these different mechanisms have qualitatively similar effects, there may be important differences between them. Finally, while we examined each scenario separately, many of them are likely to operate simultaneously, which would make archaeological evidence very challenging to interpret. We hope, however, that this study illustrates the importance of studying the dynamics that shape populations’ resilience and helps in highlighting the kinds of evidence that are relevant for distinguishing between them.

## Acknowledgements

We thank Yonatan Goldsmith, Uri Davidovich, Hai Ashkenazi, Ido Wachtel, Erella Hovers, and the Kolodny lab members for insightful discussions on these topics. This study has been supported by the Center for Sustainability at the Hebrew University of Jerusalem, the Israel Science Foundation (ISF; 1826/20), the United States – Israel Binational Science Foundation (BSF), and the Minerva Center for the study of Population Fragmentation.

